# Integrating Genetic Domestication and Model-Guided Kill-Switch Engineering in a Non-Model Bacterial Therapeutic

**DOI:** 10.64898/2026.10.18.759933

**Authors:** Yeonhwee Jeong, Hari R. Namboothiri, Oluwatola Femi-Olatunji, Paul de Figueiredo, Cansu Agca, Chelsea Y. Hu

**Affiliations:** Department of Chemical Engineering, Texas A&M University, College Station, TX 77843, USA; Bond Life Sciences Center, University of Missouri, Columbia, MO 65211, USA; Department of Molecular Microbiology and Immunology, University of Missouri School of Medicine, Columbia, MO 65212, USA; Department of Veterinary Pathobiology, University of Missouri, Columbia, MO 65211, USA; Department of Chemical and Biomedical Engineering, University of Missouri, Columbia, MO 65211, USA

**Keywords:** Genetic domestication, non-model bacteria, *Brucella melitensis* Δ*vjbR*, synthetic biology toolbox, model-guided circuit design, CRISPR-Cas9 kill switch

## Abstract

Engineered bacteria offer programmable cancer immunotherapies, but engineering tractability does not always align with therapeutic suitability. The attenuated intracellular vaccine strain *Brucella melitensis* Δ*vjbR* (BmΔ*vjbR*) shows promise as an immunotherapy chassis, yet its translation requires reliable control and containment systems that remain difficult to engineer with the limited genetic tools available. Here, we establish a genetic domestication workflow that connects genetic part generation and quantitative evaluation with model-guided circuit design. Through mutagenesis and screening, we generated and characterized genetic part libraries, including promoters spanning two orders of magnitude in activity, a panel of ribosome-binding sites, and highly functional terminators. We also evaluated fluorescent reporters, plasmid compatibility, and three inducible regulatory systems. These host-specific measurements informed mathematical modeling to guide the selection of a doxycycline-inducible Cas9/gRNA kill-switch architecture before construction. Experimental validation demonstrated inducible control both *in vitro* and *in vivo*: doxycycline reduced viable BmΔ*vjbR* below the assay detection limit *in vitro* and decreased bacterial burden in mouse liver by approximately 91%. To our knowledge, this work establishes the first systematic genetic domestication of BmΔ*vjbR* and the first containment circuit in this strain. The resulting workflow provides an extensible framework for domesticating and engineering other non-model bacteria.

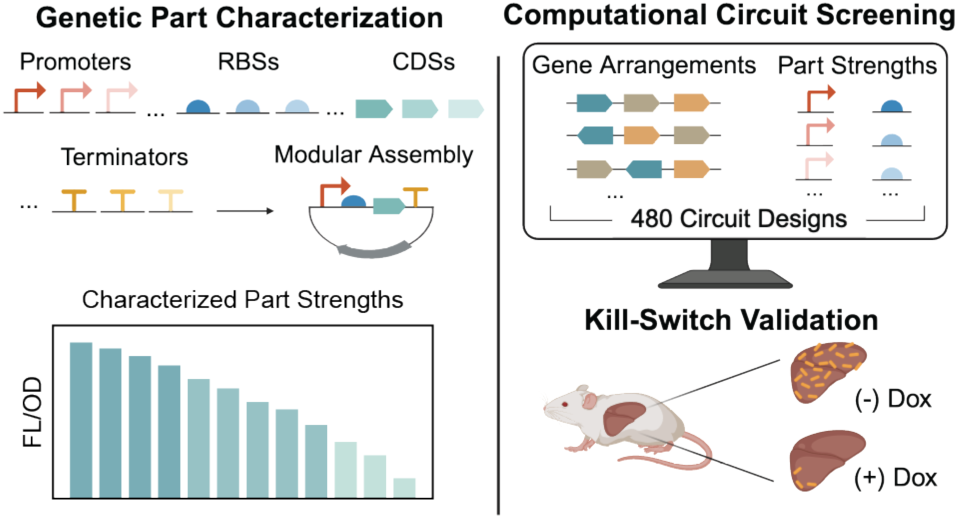

## Introduction

Bacteria are involved in nearly every aspect of human health, creating opportunities to use living bacterial cells as therapeutic agents. Recent work has demonstrated engineered bacteria for gut metabolic disease, intestinal infection, inflammation, and cancer immunotherapy.^1–3^ These studies include engineered probiotic strains that consume disease-associated metabolites,^4, 5^ sense and suppress enteric pathogens,^6^ remodel the tumor microenvironment,^3^ deliver immunostimulatory payloads,^1, 2, 7, 8^ or function as *in situ* vaccine platforms.^9, 10^ Together, these examples show that bacterial therapeutics are an active and rapidly expanding area of synthetic biology.^11^ However, most current platforms rely on a small number of genetically tractable chassis, especially *E. coli* and attenuated *Salmonella* strains.^12–16^ Although these strains benefit from established genetic engineering tools, their endotoxin-associated inflammatory activity can complicate systemic administration, where antitumor immune stimulation must be balanced against inflammatory toxicity.^7, 17, 18^ This tension motivates the exploration of alternative chassis that combine therapeutic suitability with engineering tractability.

BmΔ*vjbR* is an attenuated intracellular bacterial strain originally developed as a live vaccine candidate for brucellosis.^19^ Early studies established its safety profile in relevant animal models, including pregnant goats.^20–22^ More recently, BmΔ*vjbR* has emerged as a candidate chassis for bacterial cancer immunotherapy because it combines several properties that are desirable for systemic therapeutic delivery.^23, 24^ Unlike many commonly engineered Gram-negative probiotic strains, BmΔ*vjbR* lacks canonical endotoxin activity,^25^ reducing one major source of inflammatory toxicity. Its intracellular lifestyle provides access to immune-cell-rich tumor environments, while low pre-existing adaptive immunity in the U.S. population supports feasibility of repeated or sustained therapeutic activity.^26^ In mouse tumor models, BmΔ*vjbR* has shown strong tumor-homing capabilities, further supporting its potential as a living therapeutic chassis.^19^

Further translation of BmΔ*vjbR* requires control and containment systems to support safe use in humans. For systemically administered living therapeutics, fail-safe mechanisms are essential to restrict or eliminate engineered bacteria when their activity must be terminated.^27, 28^ Implementing such mechanisms in BmΔ*vjbR* requires quantitatively designed gene circuits built from well-characterized genetic parts. Although previous studies have demonstrated that BmΔ*vjbR* can be genetically modified,^29, 30^ genetic accessibility alone does not establish a basis for quantitative circuit design. A central challenge is connecting host-specific measurements of genetic part behavior to the circuit-level functions required for therapeutic control.

Recent efforts to domesticate non-model bacteria have established genetic parts, assembly workflows, and genome-engineering strategies across diverse chassis.^31–41^ These studies highlight the importance of characterizing parts in their intended host, where their behavior may differ from that observed in model organisms. For quantitative engineering, however, characterization must do more than establish which parts function: it must generate information that informs circuit design. Linking these measurements to mathematical models provides a framework for evaluating design choices before construction and relating component behavior to circuit-level outcomes. From this perspective, genetic domestication provides the experimental foundation for model-guided engineering.

Here, we establish a genetic domestication workflow that integrates genetic part diversification, quantitative characterization, and model-guided circuit design in BmΔ*vjbR*. We generated genetic part libraries through mutagenesis and quantitatively evaluated the resulting variants, complementing these libraries with characterized fluorescent reporters, compatible plasmid backbones, and inducible regulatory systems. We then used these measurements to inform mathematical modeling and guide the selection of a doxycycline-inducible Cas9/gRNA kill-switch architecture before construction. Experimental validation demonstrated kill-switch function both *in vitro* and *in vivo*: doxycycline induction reduced viable bacteria below the assay detection limit *in vitro* and decreased bacterial burden in mouse liver by approximately 91%. To our knowledge, this study represents the first systematic genetic domestication of BmΔ*vjbR* and the first demonstration of an engineered safety-control circuit in this strain. By connecting part generation and quantitative characterization to model-guided design and experimental validation, this work provides a framework for engineering other non-model bacterial chassis.

## Results

### Establishing recombinant gene expression in plasmid vectors

To establish BmΔ*vjbR* as an engineerable chassis for living bacterial immunotherapy, we first evaluated its capacity for heterologous gene expression using fluorescent reporters. We tested a panel of fluorescent proteins that ranged from blue to red to identify reporters suitable for quantitative characterization of gene expression in BmΔ*vjbR*. All fluorescent protein coding sequences were codon-optimized for *B. melitensis* based on the codon usage table retrieved from the Kazusa Codon Usage Database.^42^ BFP,^43^ sfCFP,^44^ eGFP,^45^ sfYFP,^46^ mCherry,^47^ and mRFP^48^ were all detectably expressed in BmΔ*vjbR*, although their signal relative to cellular background varied (Fig. 1a). Consistent with bacterial autofluorescence profiles observed in other Gram-negative bacteria, BmΔ*vjbR* exhibited elevated autofluorescence in the blue and green fluorescence microscopy channels. These results establish the relative performance of fluorescent reporters spanning multiple spectral channels and expression ranges for subsequent circuit characterization.

**Figure 1.**
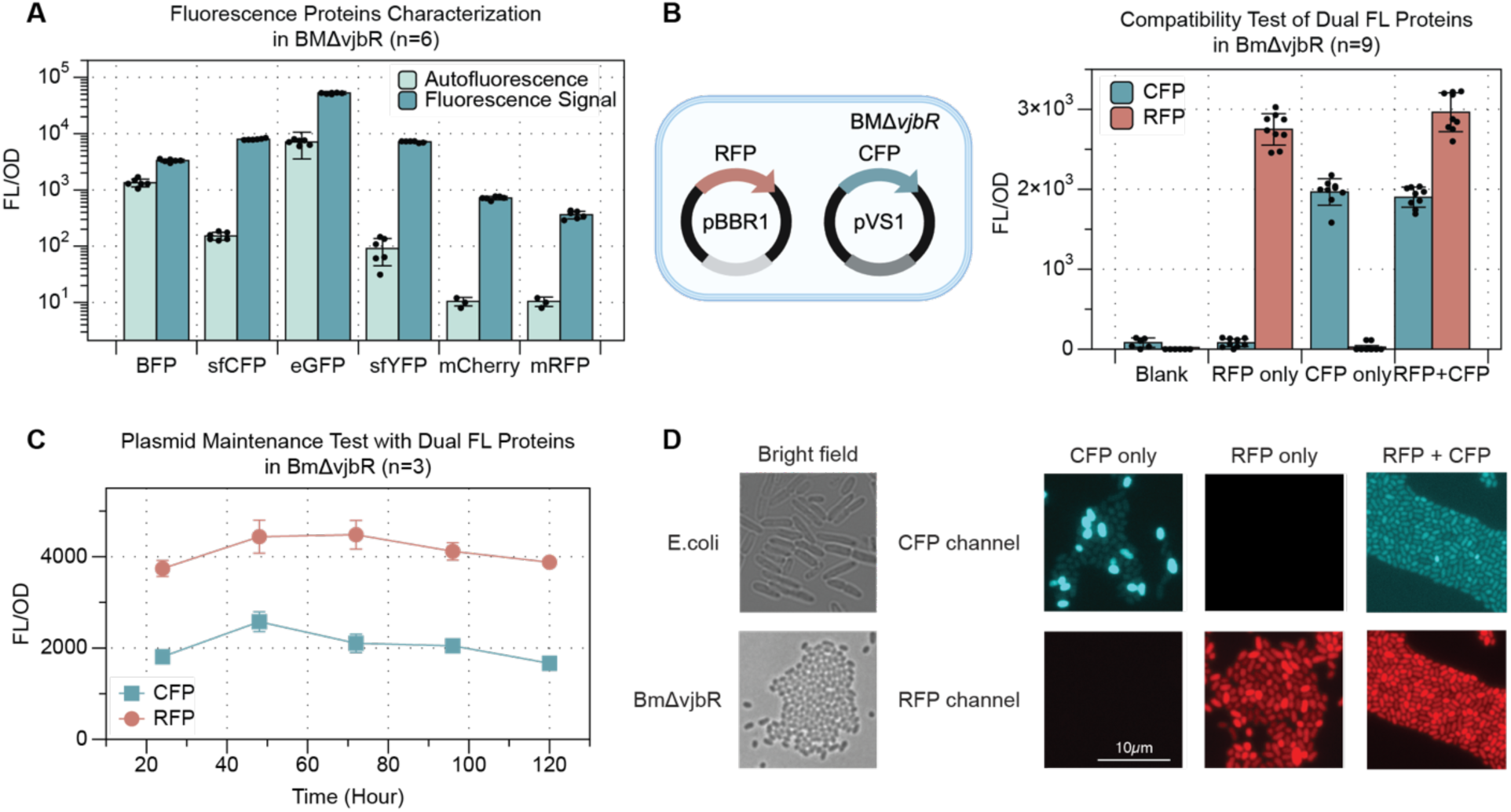
Establishing recombinant gene expression in plasmid vectors. (A) Characterization of fluorescent proteins in BmΔ*vjbR*. Autofluorescence and fluorescence signals were measured for BFP, sfCFP, eGFP, sfYFP, mCherry, and mRFP (n = 6 biological replicates). (B) Compatibility of two plasmid backbones in BmΔ*vjbR* cells co-transformed with pVS1-CFP and pBBR1-RFP was analyzed for CFP and RFP fluorescence to assess simultaneous expression from both plasmids (n = 9 biological replicates). (C) Assessment of plasmid maintenance in BmΔ*vjbR* by monitoring CFP and RFP fluorescence for 120 h (n = 3 biological replicates). (D) Representative microscopy images of *E. coli* and BmΔ*vjbR* expressing CFP and/or RFP, shown in bright-field, CFP, RFP, and merged fluorescence channels. Scale bar: 10 μm. Data are presented as mean ± SD. The schematic in Figure 1B was created using BioRender. Jeong, Y. (2027) https://BioRender.com/1tyhqgr

Because more complex genetic programs often require distinct sensing, regulatory, and output modules that can be distributed across multiple plasmids, we next tested plasmid compatibility in BmΔ*vjbR* using two fluorescent reporter plasmids. In this setup, one plasmid encoded CFP with a pVS1^49^ origin and the other encoded RFP with a pBBR1^50^ origin, allowing us to determine whether both plasmids could be maintained and expressed in the same cell. Cells carrying both plasmids exhibited detectable fluorescence from both reporters, whereas cells carrying a single plasmid produced fluorescence only from the corresponding reporter (Fig. 1b). We then monitored the dual-plasmid strain over 120 h under antibiotic selection and found that both CFP and RFP signals remained relatively stable over 5 days (Fig. 1c). These results support the compatibility of the tested plasmid pair and establish a basis for multi-component genetic circuit construction in BmΔ*vjbR*.

Finally, because BmΔ*vjbR* cells are substantially smaller than *E. coli*, with dimensions of approximately 0.6-1.5 μm in length and 0.5-0.7 μm in width,^51^ compared with ∼ 2.5 μm in length and ∼ 0.9 μm in width for *E. coli*^52^ (Fig. 1d). Because of their small size, resolving individual cells by fluorescence microscopy was technically challenging. We therefore established an agarose pad-based imaging workflow for BmΔ*vjbR* microscopy (see Methods). This approach enabled high-resolution single-cell imaging with well-separated cells and minimal overlap. Consistent with the bulk fluorescence measurements, CFP and RFP signals remained independently resolvable at the single-cell level, confirming that dual-color readout can be measured in BmΔ*vjbR* using both population-level and microscopy-based assays.

### Creating a toolbox for gene expression fine-tuning

To develop a gene-expression tuning toolbox for BmΔ*vjbR*, we first sought to identify compact constitutive promoters that could be incorporated into a modular cloning framework. We identified two *Brucella* native promoter sequences, pGroE^53^ and pTrcD,^54^ that drove measurable reporter expression in BmΔ*vjbR* (Fig. 2a, b). The reported promoter fragments included uncharacterized sequences downstream of the TSS that likely contained RBSs. To generate modular promoter parts and enable independent tuning of transcription and translation, we truncated these downstream sequences to produce minimal promoter variants, then paired each variant with a standardized MoClo-compatible RBS and showed that both minimal promoter variants remained functional (Fig. 2a, b). The minimal pTrcD promoter drove approximately 5-fold higher reporter expression than minimal pGroE, indicating distinct promoter activities under the same RBS context. These results establish pGroE and pTrcD as compact constitutive promoter parts that can be combined with interchangeable RBS sequences for modular gene-expression tuning in BmΔ*vjbR*.

**Figure 2.**
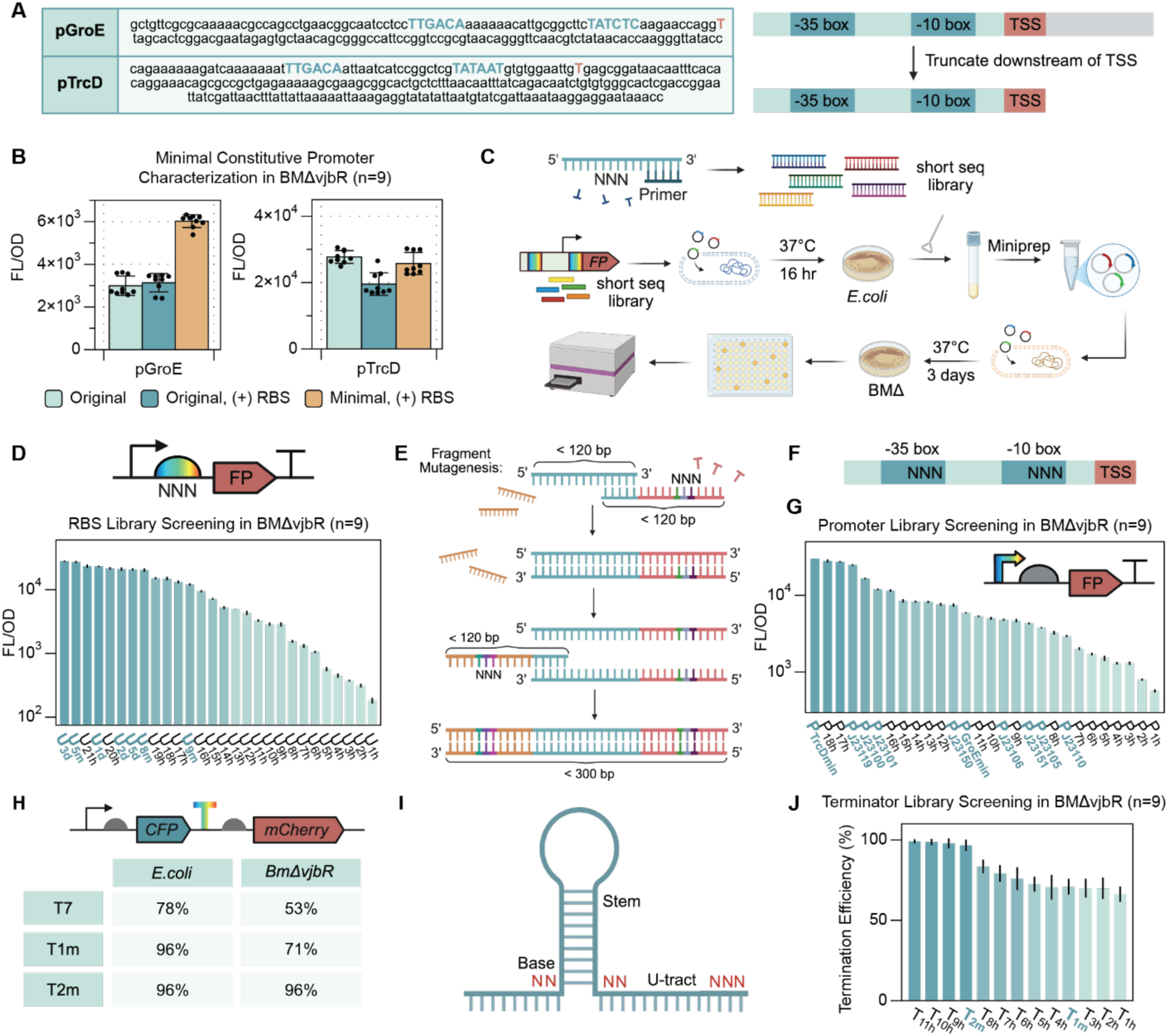
Creating a toolbox for gene expression fine-tuning. (A) Identification and truncation of the native pGroE and pTrcD promoters, showing the −35 and −10 elements, transcription start site (TSS), and uncharacterized sequences downstream of the TSS. Minimal promoter variants were generated by truncating these downstream sequences. (B) Characterization of pGroE and pTrcD promoter variants in BmΔ*vjbR*. Fluorescence output was measured for three variants of each promoter: native, native + RBS, and minimal + RBS (n = 9 biological replicates). (C) Workflow for the systematic construction and screening of short-sequence genetic part libraries. Libraries were assembled in *E. coli* and pooled after transformation. Pooled plasmids were then purified, transformed into BmΔ*vjbR*, and screened for reporter expression. (D) Fluorescence signals in BmΔ*vjbR* expressing an FP under the control of different RBS variants identified through library screening (n = 9 biological replicates). (E) Fragment mutagenesis strategy for constructing promoter libraries from three overlapping DNA fragments, each <120 bp in length. (F) Schematic of promoter diversification by site-directed mutagenesis of the −35 and −10 regions. (G) Fluorescence signals in BmΔ*vjbR* expressing a fluorescent protein under the control of promoter variants identified through library screening (n = 9 biological replicates). (H) Termination efficiencies of the T7, T1m, and T2m terminators in *E. coli* and BmΔ*vjbR*, measured using a dual-reporter assay. (I) Schematic of terminator library design, showing mutations in the stem and downstream U-rich tract. (J) Termination efficiencies of variants identified through library screening (n = 9 biological replicates). Data are presented as mean ± SD. Illustrations in Figure 2 were partially created using BioRender. Jeong, Y. (2027) https://BioRender.com/zd9×16b

We next developed a general workflow for constructing and screening expression-tuning libraries in BmΔ*vjbR* (Fig. 2c). Short sequence libraries were first assembled by Golden Gate cloning in *E. coli.*^55^ Rather than screening individual *E. coli* colonies, we pooled colonies directly from the transformation plate, isolated the mixed plasmid library by DNA extraction, and transformed the resulting library into BmΔ*vjbR*. Individual BmΔ*vjbR* colonies were then picked and screened for reporter expression. Using this workflow, we first constructed an RBS library under a defined promoter context (Supplementary Table 1). As schematized in Fig. 2b, the minimal pTrcD promoter identified above was placed upstream of an RBS library and an mCherry reporter in a single Golden Gate assembly reaction. Screening individual BmΔ*vjbR* transformants revealed a panel of RBS variants that drove mCherry expression across approximately two orders of magnitude (Fig. 2d). This RBS library therefore provides a modular set of translation-control elements for tuning recombinant protein expression in BmΔ*vjbR*.

Promoter libraries required a different construction strategy because promoter parts typically have lengths beyond the practical range of the ∼120-nt oligonucleotides used for two-primer annealing. We therefore used fragment mutagenesis to generate MoClo-compatible promoter libraries up to ∼300 nt (Fig. 2e, f). Three primers, each shorter than 120 bp and containing 18-24 nt overlapping regions, were assembled into full promoter variants while preserving the flanking MoClo adapters (Supplementary Table 2). For each promoter template, we applied site-directed mutagenesis to the −35 and −10 regions. This allows us to diversify expression strength by perturbing the core promoter elements. Using minimal pTrcD and pGroE as templates, we pooled the assembled libraries through *E. coli*, transformed the recovered plasmid library into BmΔ*vjbR,* and screened individual BmΔ*vjbR* colonies as described above. This yielded promoter variants that tuned reporter expression across approximately two orders of magnitude (Fig. 2g). We also characterized eight Anderson constitutive promoters^56^ commonly used in *E. coli* (highlighted in green in Fig. 1g) and incorporated them into the promoter set, expanding the library to include both native promoter-derived variants and established bacterial constitutive promoters.

Because terminator efficiency can depend on host transcription machinery and sequence context, terminators characterized in model organisms may not perform equivalently in BmΔ*vjbR*. We therefore evaluated three commonly used *E. coli* terminators frequently deployed in gene circuit construction. Following the terminator characterization and library-screening framework described by Ren et al.,^57^ terminator performance was measured using a dual-reporter circuit in which CFP and mCherry, previously characterized in Fig. 1, were separated by the candidate terminator sequence (Fig. 2h).^58^ In this design, readthrough transcription produces downstream mCherry expression, so a higher mCherry/CFP ratio indicates weaker termination. Terminator efficiency was calculated from this ratio as described in the Methods. Two of the three tested *E. coli* terminators showed poor termination efficiency in BmΔ*vjbR*, motivating the construction of a host-validated terminator library. Using the same framework, we generated this library from two parental terminators, T1m and T2m, by introducing mutations into the stem sequence and the downstream U-rich tract (Supplementary Table 3).^57^ The resulting libraries were constructed and screened using the same pooled-library workflow described in Fig. 2c. This screen identified terminator variants with graded termination efficiencies, including variants reaching up to 99% efficiency in BmΔ*vjbR* (Fig. 2j). Together with the promoter and RBS libraries, these results establish a modular expression-tuning toolbox for controlling transcriptional, translational, and transcription-termination strength in BmΔ*vjbR*.

### Establishing inducible gene regulation in BmΔ*vjbR*

To extend the regulatory toolbox of BmΔ*vjbR* beyond constitutive expression, we tested three inducible systems that respond to distinct inputs: LacI/IPTG, TetR/aTc, and the temperature-responsive TlpA module.^59, 60^ Each circuit was constructed using a common architecture: the regulator was expressed constitutively, while mCherry was placed downstream of the cognate inducible promoter. We quantified each circuit by induction strength, dose response, basal expression, and growth impact, with the goal of identifying inducible modules that could support tunable gene expression during strain construction, circuit characterization, and future therapeutic programming in BmΔ*vjbR*.

The LacI/IPTG system was evaluated as a chemically inducible module in BmΔ*vjbR*. LacI was constitutively expressed, while mCherry was placed under the LacI-repressed pLac promoter (Fig. 3a). IPTG induced a dose-dependent increase in mCherry expression, reaching approximately 170–180-fold activation before plateauing (Fig. 3b). Hill-function fitting yielded an apparent induction constant of ∼0.105 mM IPTG and a Hill coefficient of ∼1.46. Growth was largely unaffected across most IPTG concentrations, with reduced OD_600_ only at the highest dose tested (Fig. 3c). Thus, LacI/IPTG provides strong inducible expression in BmΔ*vjbR*, although high IPTG concentrations may impose a modest growth cost.

**Figure 3.**
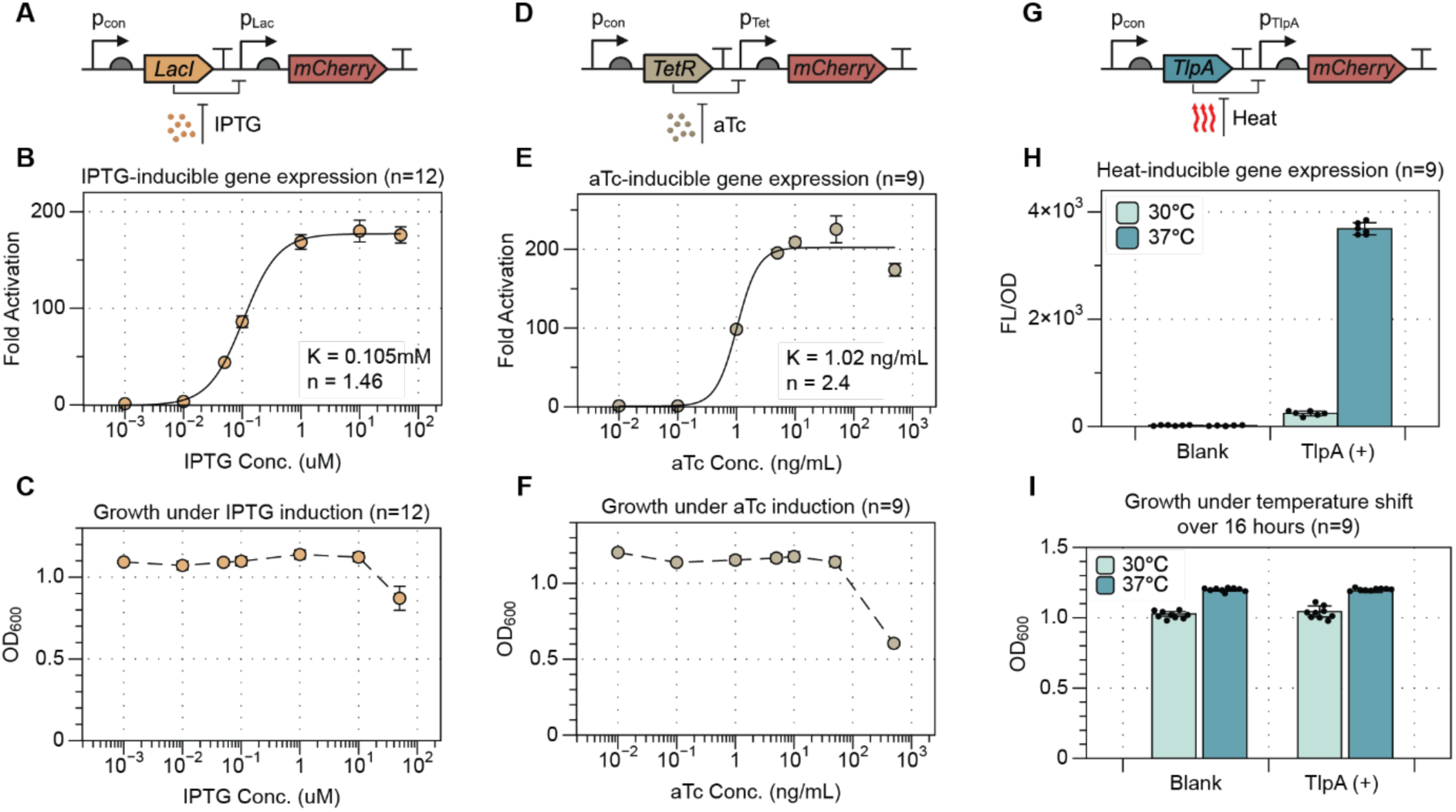
Characterization of inducible regulatory systems in BMΔ*vjbR*. (a) Schematic of the IPTG-inducible expression circuit, in which LacI represses the pLac promoter controlling mCherry expression. (b) Dose–response characterization of IPTG-inducible gene expression in BmΔ*vjbR*. Fold activation was quantified across increasing IPTG concentrations and fitted to a Hill function, yielding K = 0.105 mM and n = 1.46 (n = 9 biological replicates). (c) OD_600_ of BmΔ*vjbR* after 16 h under increasing IPTG concentrations, measured by OD600 (n = 9 biological replicates). (d) Schematic of the aTc-inducible expression circuit, in which TetR represses the pTet promoter controlling mCherry expression. (e) Dose–response characterization of aTc-inducible gene expression in BmΔ*vjbR*. Fold activation was quantified across increasing aTc concentrations and fitted to a Hill function, yielding K = 1.02 ng/mL and n = 2.4 (n = 9 biological). (f) OD_600_ of BmΔ*vjbR* after 16 h under increasing aTc concentrations, measured by OD600 (n = 9 biological replicates). (g) Schematic of the temperature-responsive TlpA expression circuit controlling mCherry expression. (h) Temperature-dependent reporter expression in BmΔ*vjbR* carrying the TlpA regulatory system at 30 °C and 37 °C (n = 9 biological replicates). (i) OD_600_ of BmΔ*vjbR* after 16 h in control and TlpA-containing BmΔ*vjbR* cultures at 30 °C and 37 °C (n = 9 biological replicates). Data are presented as mean ± SD. Illustrations in Figure 3 were created using BioRender. Jeong, Y. (2027) https://BioRender.com/01e4m6w

We next evaluated the TetR/aTc system as a second chemically inducible module in BmΔ*vjbR*. TetR was constitutively expressed, while mCherry was placed under the TetR-repressed pTet promoter (Fig. 3d). aTc induced a dose-dependent increase in fluorescence, reaching >200-fold activation before declining at the highest inducer concentration (Fig. 3e). Hill-function fitting yielded an apparent induction constant of ∼ 1.02 ng/mL aTc and a Hill coefficient of ∼ 2.4. Growth was largely maintained across the induction range but decreased at the highest aTc concentration, coinciding with reduced fluorescence (Fig. 3f). Thus, TetR/aTc provides high dynamic range inducible expression in BmΔ*vjbR*, but excessive aTc induction can impair growth and reduce circuit output.

To incorporate non-chemical regulation, we next tested the temperature-sensitive TlpA system, in which increased temperature relieves TlpA-mediated repression. mCherry expression remained near background after growth at 30°C for 16 h but was strongly activated after shifting to 37°C, producing a clear OFF/ON transition between the two conditions (Fig. 3g, h). OD_600_ measurements showed higher growth at 37°C than at 30°C, indicating that circuit output is coupled to temperature-dependent host growth (Fig. 3i). Thus, TlpA provides effective heat-inducible switching in BmΔ*vjbR*, although applications using this system should account for reduced growth under the lower-temperature OFF condition.

Together, these results establish LacI/IPTG, TetR/aTc, and TlpA as functional inducible systems in BmΔ*vjbR*. The two chemical systems provide dose-dependent control with high dynamic range, while TlpA adds a temperature-based input. These data define a set of validated inducible modules for tuning gene expression in BmΔ*vjbR* and provide the regulatory foundation for constructing more complex therapeutic circuits and safety-control systems.

### Modeled guided design of doxycycline-inducible kill-switch in BmΔ*vjbR*

Building on the expression parts and inducible systems established above, we next applied this toolbox to design a programmable kill-switch for BmΔ*vjbR*. For living bacterial therapeutics, controllable elimination of the engineered strain is an important safety feature, particularly if treatment must be stopped after administration. Building on the CRISPR-based biocontainment strategy reported by Rottinghaus et al.,^61^ we designed a doxycycline-inducible Cas9 kill-switch in which induction activates expression of a gRNA and Cas9 targeting the BmΔ*vjbR* genome.^62, 63^ In the OFF state, TetR represses pTet and inhibits expression of the lethal module. In the ON state, doxycycline relieves TetR repression, allowing Cas9–gRNA complex formation and genome cleavage.

Although the required parts were available, assembling them into an effective kill-switch was not straightforward. The circuit needed to satisfy two competing requirements: low basal Cas9–gRNA activity in the absence of inducer, and high induced activity after doxycycline addition. Because TetR, Cas9, and the gRNA could be arranged in different transcriptional orientations and assigned different promoters and RBSs, we performed a computational screen of 480 possible circuit variants (Fig. 4a). The screen varied module order, orientation, promoter assignment, and RBS strength, allowing us to evaluate a broad but experimentally tractable design space.

**Figure 4.**
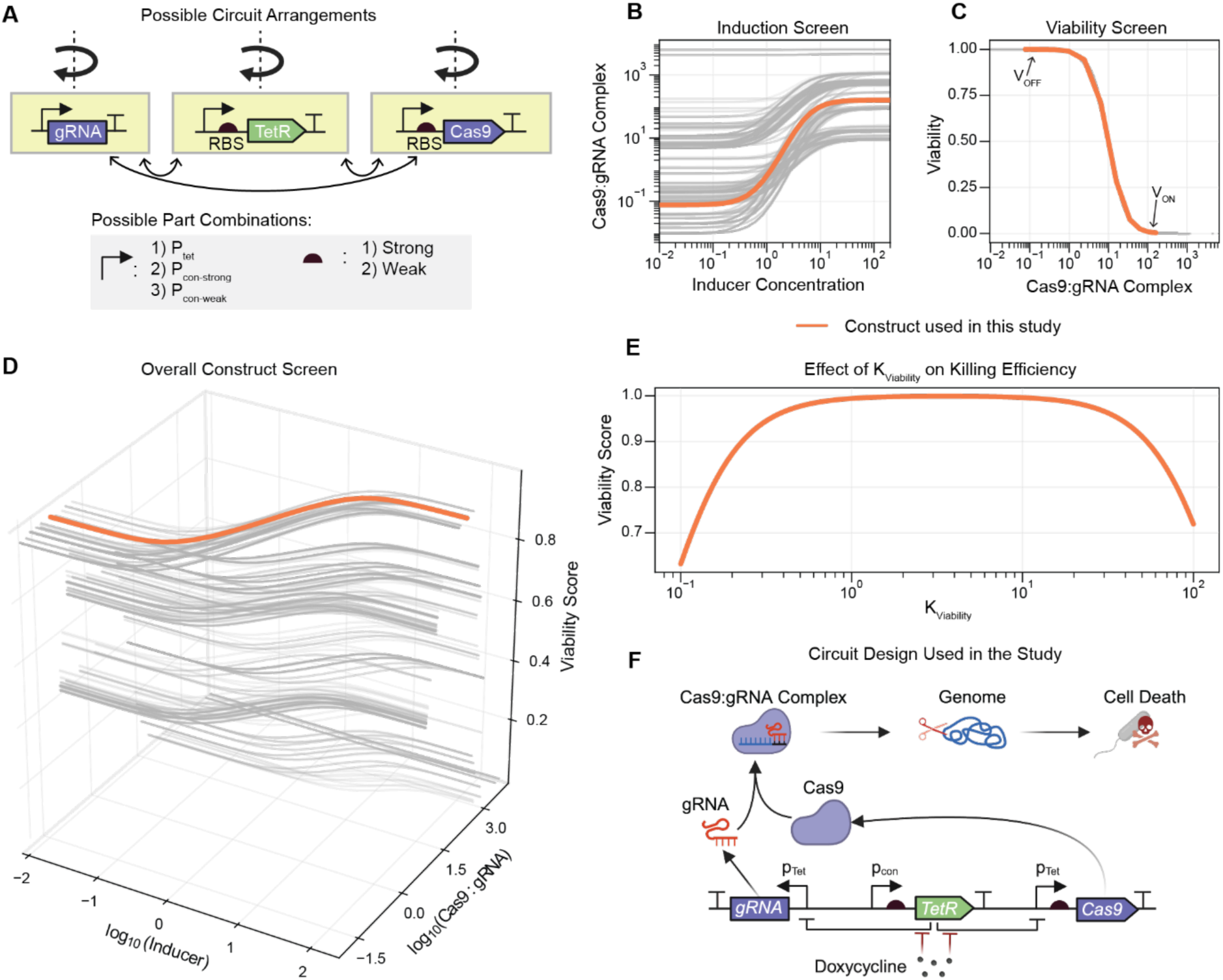
Modeled guided design of a doxycycline-inducible kill-switch in BMΔ*vjbR*. (A) Schematic of the kill-switch design space, showing alternative arrangements of the gRNA, TetR, and Cas9 expression cassettes. Gene arrangement, promoter strength, and RBS strength were varied to generate circuit configurations for subsequent screening and optimization. (B) Induction screen showing the dose-response curves for all circuit combinations. The y-axis is Cas9-gRNA complex concentration, and the x-axis is inducer concentration. (C) Viability screen showing viability as a function of Cas9-gRNA complex concentration, using the concentration range defined by the y-axis of the induction screen. (D) 3D plot showing how the viability score changes with inducer concentration and Cas9-gRNA complex concentration. The highlighted line represents the circuit used in the study. (E) Viability score as a function of K_via_, where K_via_ is linked to the number of genomic target sites for cutting. (F) Kill-switch schematic reflecting the circuit topology selected to proceed in this study. Illustrations in Figure 4 were created using BioRender. Jeong, Y. (2027) https://BioRender.com/z1990ez

For each circuit variant, we first simulated the doxycycline dose-response and calculated the steady-state concentration of active Cas9:gRNA complex (see Supplementary Method 2 for the model description), denoted *C_Cas_*_$:*sRNA*_ (Fig. 4b). This quantity represents the effective abundance of the lethal species produced by each circuit at a given doxycycline concentration. Designs with high basal *C_Cas_*_$:*sRNA*_ would be expected to reduce cell viability even in the OFF state, whereas designs with insufficient induced *C_Cas_*_$:*sRNA*_ would fail to kill efficiently in the ON state.

We next converted predicted Cas9:gRNA abundance into predicted cell viability using a negative Hill-type function:

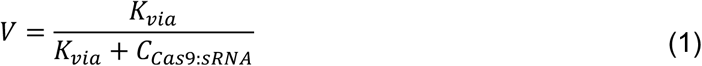

where *V* is the predicted cell viability, *C_Cas_*_$:*sRNA*_ is the active Cas9–gRNA complex concentration, and *K_via_* is the Cas9:gRNA complex concentration at which viability is reduced by 50% (Fig. 4c). Thus, *K_via_* represents the effective killing sensitivity of the strain to Cas9–gRNA activity. A lower *K_via_* indicates greater sensitivity, whereas a higher *K_via_* indicates that more Cas9:gRNA complex is required to reduce viability.

For each circuit, we used this function to calculate predicted viability in the OFF and ON states. We define *V*_0*FF*_ as the predicted viability without doxycycline, and *V*_0*N*_ as the predicted viability under inducing conditions. We then ranked circuit variants using a viability score *S_via_* that rewards high OFF-state viability and low ON-state viability:

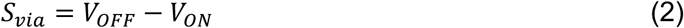

In this formulation, high-scoring circuits preserve viability without inducer while producing strong killing after induction. This score was used to compare all circuit variants across the simulated design space (Fig. 4d). The model identified designs with a range of predicted behaviors, including leaky circuits with reduced OFF-state viability, weakly induced circuits with poor ON-state killing, and switch-like circuits with favorable OFF/ON separation.

We then examined how the assumed killing sensitivity affected circuit performance. For a high-scoring architecture, we varied the effective killing sensitivity, *K_via_*, defined in Eq. (1), and recalculated *S_via_* (Fig. 4e). The score was maximal at intermediate *K_via_* values and decreased when *K_via_* was either too high or too low. When *K_via_* is high, the maximum concentration of Cas9–gRNA complex is not sufficient to drive efficient killing. When *K_via_* is too low, basal Cas9– gRNA expression can reduce OFF-state viability. This analysis indicated that optimal kill-switch behavior requires an intermediate effective killing sensitivity.

This killing sensitivity analysis informed gRNA target selection. Candidate gRNAs were identified from valid PAM-containing genomic sites and prioritized based on target copy number, predicted on-target activity, and off-target specificity (see Methods). Because the objective was killing rather than genome editing, target sequences present at multiple genomic loci were preferred, as they could generate multiple double-strand breaks after induction. One candidate targeting six genomic loci while retaining the highest predicted on-target activity was selected.

The final model-guided architecture is shown in Fig. 4f and highlighted in orange in the computational screen (Fig. 4b–d). In this design, both the gRNA and Cas9 are controlled by pTet. TetR is expressed from a constitutive promoter with a medium-strength RBS, while Cas9 is paired with a weak RBS to reduce OFF-state activity. This architecture was selected to balance low basal toxicity with strong induced killing while remaining experimentally tractable. We next constructed this circuit and evaluated its kill-switch activity in BmΔ*vjbR*.

### *In vitro* and *in vivo* evaluation of the kill-switch in BmΔ*vjbR*

After selecting the model-guided kill-switch architecture, we constructed the circuit in BmΔ*vjbR* and first tested doxycycline-dependent killing *in vitro* using a CFU-based assay. The kill-switch plasmid was transformed into BmΔ*vjbR*, individual colonies were recovered, grown in liquid culture, induced with doxycycline, serially diluted, and plated to quantify viable cells (Fig. 5a). In the absence of doxycycline, the kill-switch strain produced CFUs comparable to parental BmΔ*vjbR*, indicating limited basal toxicity from the circuit. Upon doxycycline induction, recoverable CFUs decreased from approximately 10^9^ CFU/mL to below the limit of detection in the kill-switch strain, whereas parental BmΔ*vjbR* remained viable under the same condition (Fig. 5b). This result showed that activation of the pTet-controlled Cas9/gRNA module eliminated recoverable viable cells under the tested *in vitro* condition.

**Figure 5.**
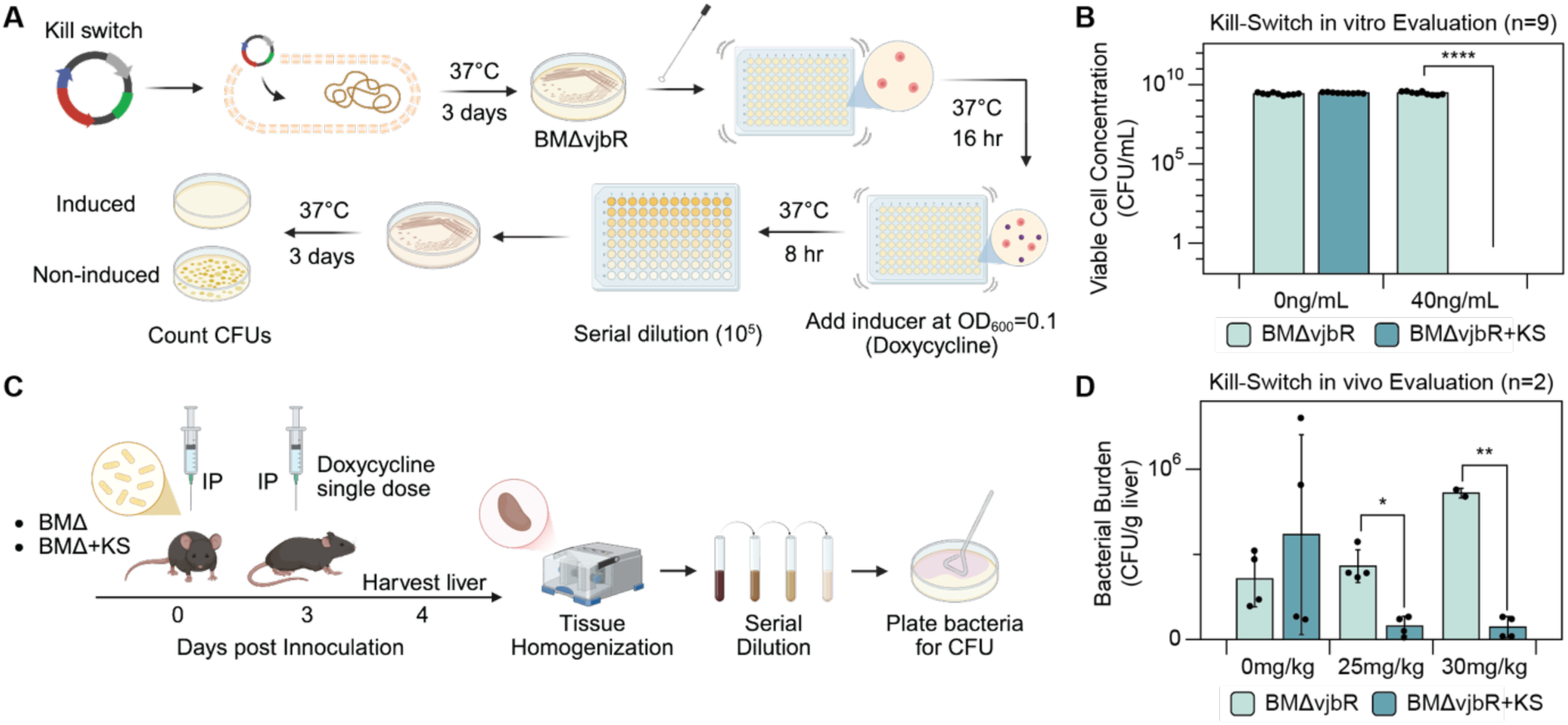
*In vitro* and *in vivo* evaluation of the kill-switch in BmΔ*vjbR*. (A) Schematic of the *in vitro* kill-switch assay. BmΔ*vjbR* carrying the kill-switch construct was cultured, induced with doxycycline, serially diluted, and plated for colony-forming units (CFUs). (B) *In vitro* kill-switch activity following doxycycline induction. CFUs were quantified for BmΔ*vjbR* with or without the kill-switch construct in the absence or presence of 40 ng/mL doxycycline (n = 9 biological replicates). (C) Schematic of the *in vivo* kill-switch assay. C57BL/6 mice were intraperitoneally inoculated with 1.07×10^9^ CFU of either BmΔ*vjbR* or BmΔ*vjbR* with the kill-switch construct. Three days after inoculation, mice received a single intraperitoneal dose of doxycycline, and livers were harvested 24 h later for tissue homogenization, serial dilution, and CFU counting. (D) *In vivo* bacterial recovery from liver following doxycycline treatment at 0, 25, or 30 mg/kg (n = 2 mice per group). Data are presented as mean ± SD. Statistical significance was assessed using two-tailed Welch’s t-tests; P < 0.05, *P < 0.01, **P < 0.001, and ***P < 0.0001. Illustrations in Figure 5 were created using BioRender. Jeong, Y. (2027) https://BioRender.com/y7rz74b

We next evaluated whether the kill-switch could reduce bacterial burden *in vivo.*C57BL/6 mice were inoculated intraperitoneally with either parental BmΔ*vjbR* or the kill-switch strain, treated with a single doxycycline dose three days after inoculation, and euthanized one day later for liver harvest and CFU quantification (Fig. 5c). In the absence of doxycycline, the kill-switch strain was recovered at levels comparable to parental BmΔ*vjbR*, consistent with limited OFF-state activity *in vivo*. Doxycycline treatment reduced recoverable CFUs from the kill-switch strain, with significant reductions observed at 25 and 30 mg/kg compared with parental BmΔ*vjbR* under the same dosing conditions (Fig. 5d). At a doxycycline dose of 30 mg/kg, the kill-switch strain showed an approximately 91% reduction in recoverable bacterial burden relative to the corresponding parental control. These data show that the model-guided circuit enables inducible reduction of BmΔ*vjbR* burden both in vitro and *in vivo*.

These results show that the engineered circuit enables doxycycline-inducible BmΔ*vjbR* cell-fate control *in vitro* and *in vivo*, providing a controllable safety mechanism for regulating persistence after administration. The *in vivo* reduction was less complete than the apparent killing observed *in vitro*, which we attribute in part to plasmid loss during animal colonization. Future versions of the kill-switch will therefore benefit from genomic integration of the circuit to improve stability and killing consistency *in vivo*.

## Discussion

This study establishes BmΔ*vjbR* as a genetically programmable chassis for living bacterial immunotherapy. Although BmΔ*vjbR* has several properties that make it attractive for therapeutic development, including intracellular tropism, attenuation, and reduced endotoxin-associated inflammatory risk,^20–25^ its use as an engineered therapeutic chassis has been limited by the lack of a validated synthetic biology toolbox. Here, we move from basic recombinant expression to expression tuning, inducible regulation, model-guided circuit design, and functional kill-switch implementation. This study establishes a systematic workflow that connects genetic part screening and quantitative characterization with model-guided circuit design in an emerging therapeutic strain.

A growing body of work has established synthetic biology toolboxes for non-model bacteria, including new assembly methods, promoter and RBS libraries, genome-editing tools, and inducible systems. These studies have made clear that genetic parts cannot simply be assumed to function identically across bacterial hosts. Promoter activity, translation initiation, transcription termination, and regulator performance can all change with host physiology and genetic context. Our results are consistent with this principle. Parts commonly used in *E. coli* required direct validation in BmΔ*vjbR*, and some elements did not retain the expected behavior. By characterizing reporters, promoters, RBSs, terminators, and inducible systems directly in BmΔ*vjbR*, this work fills a practical gap for a therapeutic strain whose biological properties are promising but whose engineering infrastructure has remained limited.

A key lesson from this study is that domestication should not stop at cataloging parts. For therapeutic engineering, the relevant question is whether characterized parts can support functional genetic programs with defined design objectives. In this work, quantitative part measurements were connected to the design of a doxycycline-inducible Cas9 kill-switch. This allowed circuit variants to be evaluated in silico before construction, reducing the number of experimentally tested architectures and providing a rational basis for selecting promoter, RBS, regulator, and gRNA configurations. This model-guided approach is particularly useful for non-model chassis engineering, where each round of cloning and testing can be slower, less predictable, and more resource-intensive than in established laboratory organisms.

The kill-switch results also illustrate an important design principle for bacterial therapeutic control: effective safety circuits must balance OFF-state stability with ON-state potency. A circuit that is too leaky can compromise strain viability before therapy is complete, while a circuit with insufficient induced expression may fail to eliminate the strain when needed. In the Cas9/gRNA system, this balance depends on inducible expression strength and the host’s sensitivity to genome cleavage, which is influenced by gRNA activity, target copy number, chromosomal accessibility, and circuit burden. By explicitly modeling this tradeoff, the design workflow identified a circuit architecture intended to preserve viability without inducer while enabling killing after doxycycline exposure.

The experimental validation supports this design logic but also reveals limitations that will be important for future development. The kill-switch produced strong doxycycline-dependent loss of recoverable cells *in vitro*, whereas the *in vivo* effect was more partial. This difference likely reflects additional constraints present in animal settings, including heterogeneous bacterial growth states, variable doxycycline exposure across tissues, immune interactions, and plasmid loss during colonization. The current kill-switch is plasmid-borne, which is useful for rapid construction and testing but not ideal for long-term therapeutic deployment. Genomic integration of the circuit will likely be required to improve stability, reduce plasmid-loss escape, and support more consistent control of bacterial persistence *in vivo*.

Additional limitations should also be considered. The toolbox developed here provides a foundation, but it does not yet define the full design space for BmΔ*vjbR* engineering. Part behavior was characterized under selected laboratory conditions, and expression levels may shift across growth phases, intracellular environments, tissues, or inflammatory states. The inducible systems tested here provide useful control inputs, but each has practical constraints, including inducer toxicity, temperature-dependent growth effects, and possible differences between culture and animal dosing. The Cas9/gRNA kill-switch provides a proof of concept for programmable elimination, but future versions will need to be evaluated for genetic stability, escape frequency, dose timing, tissue-specific efficacy, and compatibility with therapeutic payloads.

Despite these limitations, the broader significance of this work is that it provides a path for moving therapeutic bacterial strains beyond descriptive biology and into programmable engineering. Many candidate therapeutic bacteria have useful biological behaviors, such as tissue tropism, immune modulation, or niche-specific persistence, but remain difficult to engineer predictably. A rapid domestication framework that combines host-specific part validation with quantitative circuit design can help close this gap. The same logic should be adaptable to other emerging bacterial chassis, although the specific parts and performance ranges will need to be re-established in each chassis.

Together, this work establishes a synthetic biology foundation for programming BmΔ*vjbR* as a living bacterial immunotherapy chassis. By building a chassis-validated toolbox and using it to construct a functional inducible kill-switch, we show that BmΔ*vjbR* can be engineered not only to express recombinant genes, but to implement controllable cell fate decisions. Future work will focus on integrating safety circuits into the genome, coupling them with therapeutic payloads, and testing their performance in disease-relevant models. More broadly, this study supports a domestication strategy in which non-model therapeutic bacteria are rapidly equipped with the parts, measurements, and models needed for predictable therapeutic programming.

## Materials and Methods

### Reagents

NEB Turbo Competent *E. coli* (New England Biolabs, Cat. No. C2984) was used for routine plasmid cloning and propagation. An attenuated *Brucella melitensis* strain derived from strain 16M by deletion of the *vjbR* gene (*B. melitensis* 16M Δ*vjbR*; hereafter BmΔ*vjbR*) was used for functional characterization of the genetic constructs unless otherwise indicated.

The plasmids used in this study incorporated replication origins derived from pBBR1MCS-2 (Addgene #85168) and pVS1 (Addgene #187874).

### Biological resources

*E. coli* was cultured in Luria–Bertani (LB) medium (Fisher Scientific, Cat. No. BP1426-500). BmΔ*vjbR* was cultured in Brucella Broth medium (Research Products International, Cat. No. B62000-500.0), hereafter referred to as BBM, and on Brucella Broth agar, hereafter referred to as BBA. Appropriate antibiotics were added as required for plasmid selection.

The primers used in this study are listed in Supplementary Tables S1, S2, and S3. Plasmid DNA from cloning strains was prepared using the ZymoPURE Plasmid Miniprep Kits (Zymo Research, Cat. No. D4210). PCR (polymerase chain reaction) fragments were purified using the ZymoClean Gel DNA Recovery Kit (Zymo Research, Cat. No. D4007). For cloning, Q5 Hot Start DNA Polymerase, restriction enzymes, and NEBuilder HiFi DNA Assembly master mix were purchased from New England Biolabs. SOC medium (Fisher Scientific, Cat. No. 15-544-034) was used for post-transformation recovery of cells. Isopropyl β-D-1-thiogalactopyranoside (IPTG; Fisher Scientific, Cat. No. BP1755-10), anhydrotetracycline hydrochloride (aTc; Fisher Scientific, Cat. No. AAJ66688MA), and doxycycline (Fisher Scientific, Cat. No. BP26535) were used for inducible gene expression and kill-switch assay, respectively. All oligonucleotide primers used in this study are listed in Supplementary Tables S1–S3.

### Plasmid construction and transformation

All plasmids used in this study were constructed using the MoClo-based Golden Gate– Gibson (3G) assembly^55, 64, 65^. Briefly, standardized genetic parts, including promoters, ribosome-binding sites, coding sequences, and transcriptional terminators, were assembled into transcriptional units by Golden Gate assembly. The resulting transcriptional units were subsequently assembled with the appropriate plasmid backbones using Gibson assembly to generate the final constructs. Constructed plasmids were verified by plasmid sequencing.

For routine plasmid construction and propagation, assembled vectors were transformed into chemically competent *E. coli* Turbo by heat-shock transformation. Following transformation, cells were allowed to recover in SOC medium before being plated on LB agar supplemented with the appropriate selective antibiotic and incubated overnight at 37 °C. Individual colonies were subsequently cultured for plasmid propagation, and plasmid DNA was isolated and verified prior to introduction into the target bacteria.

For transformation into BmΔ*vjbR*, plasmids were introduced using a spheroplast-based electroporation method. Cells were cultured overnight in BBM following inoculation from an agar plate. The overnight cultures were then diluted 50-fold into fresh BBM supplemented with 0.2 M sucrose and 10 µg/mL ampicillin and grown overnight until they reached an OD_600_ of 0.4-0.6. During this incubation, the cell walls were partially removed under osmoprotective conditions to generate spheroplasts. The spheroplasts were gently washed and resuspended in sucrose and glycerol, mixed with purified plasmid DNA, and subjected to electroporation. Immediately after electroporation, cells were transferred to SOC medium to facilitate recovery and cell-wall regeneration and plated on BBA supplemented with the appropriate selective antibiotic and incubated for 3 days at 37 °C. Individual transformants were subsequently screened for the presence of the expected plasmid construct.

### Bulk fluorescence measurements

For screening experiments, individual colonies were inoculated into 200 μL of BBM containing antibiotic in a 200 µL 96-well plate and grown overnight at 37 °C at 1,000 rpm in a benchtop shaker. 10 µL of the overnight culture was added to 190 µL of BBM containing antibiotics and grown for 6 h at 37 °C at 1,000rpm.

For single time-point measurements, the preculture was then diluted 10-fold with BBM containing its appropriate antibiotics, then grown overnight in a black 96-well plate. Fluorescence was measured using fluorophore-specific excitation/emission wavelengths: BFP, 402/457nm; sfCFP, 433/475nm; eGFP, 488/507nm; sfYFP, 485/528nm; mRFP, 584/607nm; and mCherry, 587/610nm. Optical density was measured at 600nm (OD_600_).

### Fluorescence microscopy

To observe fluorescence signals at single-cell resolution, harvested cells were collected by centrifugation (3,500 rpm, 5 min) to obtain a cell pellet. The pellet was resuspended in melted agarose gel (1%, w/v in BBM) and immediately mounted on a cover glass to immobilize the cells for imaging.

Fluorescence imaging was performed using a motorized inverted microscope (ECLIPSE Ti2-E, Nikon, Tokyo, Japan) equipped with a digital sCMOS camera (ORCA-Fusion C14440-20UP, Hamamatsu Photonics, Hamamatsu, Japan) and a 100× oil-immersion objective lens (NA 1.45). Single-cell fluorescence signals were captured with exposure times of 70 ms for CFP and 60 ms for mCherry, using appropriate filter sets for cyan and red fluorescence, respectively. Images were acquired and analyzed using NIS-Elements (Nikon).

### Construction and screening of short-sequence genetic-part libraries

Short-sequence genetic-part libraries were generated from oligonucleotide templates containing degenerate nucleotide positions using library-specific primers. The resulting library fragments were assembled into the corresponding fluorescent reporter plasmids and transformed into *E. coli* Turbo-competent cells via heat shock. Transformants were selected on LB agar plates containing 50 µg/mL kanamycin, grown at 37 °C for 16 h, and pooled to preserve library diversity. Plasmid DNA was then extracted from the pooled *E. coli* population and transformed into BmΔ*vjbR*.

Following incubation at 37 °C for 3 days, individual BmΔvjbR transformant colonies were picked into 96-well plates containing 200 µL of BBM supplemented with 50 µg/mL kanamycin and cultured overnight at 37 °C. For subculturing, 10 µL of the overnight culture was diluted into 190 µL of fresh BBM and incubated for 6 h. The pre-culture was then diluted 10-fold into fresh BBM to initiate the assay. Following overnight incubation, cell density (OD_600_) and fluorescence intensity (mCherry; excitation: 575 nm, emission: 610 nm) were measured using a microplate reader (BioTek Synergy H1) to evaluate reporter expression.

### Quantification of transcriptional termination efficiency

Termination efficiency was quantified using a dual-fluorescence reporter consisting of an upstream CFP and a downstream mCherry reporter separated by the test terminator^57^. A terminator-free construct containing the same reporter architecture was used as the positive control. Following the same experimental procedure used for bulk fluorescence measurements, mCherry and CFP signals were measured. For each construct, downstream reporter expression was normalized to the upstream CFP signal by calculating the mCherry/CFP fluorescence ratio. The normalized readthrough of each terminator was then calculated relative to the terminator-free positive control, and termination efficiency was determined as:

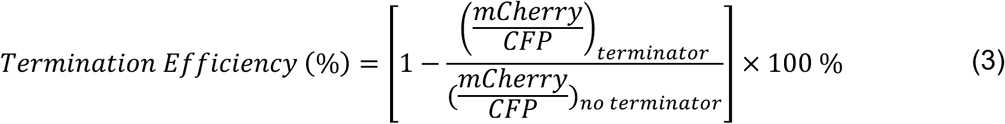

### Temperature-inducible gene expression

For characterization of temperature-inducible gene expression, which is TlpA system, individual colonies were inoculated into 200 µL of BBM containing the appropriate antibiotics in 96-well plates and grown overnight at 30 °C with shaking at 1,000 rpm. Overnight cultures were diluted by transferring 10 µL of culture into 190 µL of fresh BBM containing the appropriate antibiotics and grown for an additional 6 h at 30 °C with shaking at 1,000 rpm. Cultures were then diluted 10 times with BBM containing the appropriate antibiotics and incubated overnight at either 30 °C (non-induced condition) or 37 °C (induced condition). mCherry fluorescence (575 nm excitation, 610 nm emission) and optical density at 600 nm (OD_600_) were measured at the end of incubation. Fluorescence was normalized to OD_600_ (FL/OD) to quantify temperature-dependent gene expression.

### Inducer dose-response fitting

Inducer dose-response curves were fitted using a Hill function:

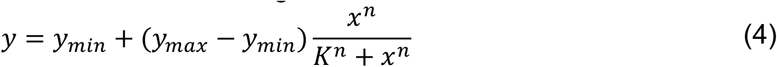

where *y* is the fold activation, *x* is the inducer concentration, *y_min_* and *y_max_* are the minimum and maximum fold activation, respectively, *K* is the inducer concentration producing half-maximal activation, and *n* is the Hill coefficient. Nonlinear least-squares fitting was performed using the GRG Nonlinear algorithm in Microsoft Excel Solver by minimizing the sum of squared residuals between the observed and predicted values. Because fold activation was normalized to the uninduced condition, *y_min_* was fixed at 1, while *K* and n were fitted.

### Computational identification of gRNA targets

Candidate Cas9 targets were identified computationally across the BMΔ*vjbR* genome by searching for genomic sequences adjacent to the canonical Cas9 protospacer-adjacent motif (PAM), NGG. All PAM-compatible target sequences were extracted, after which candidates occurring at multiple genomic locations were prioritized to enable simultaneous cleavage at repeated target sites. Candidate gRNAs were then evaluated using Benchling (https://benchling.com) for predicted on-target activity and off-target specificity. Based on these scores, we selected a gRNA targeting a sequence present at six genomic locations and exhibiting high predicted on-target activity and low predicted off-target potential. This gRNA was used for subsequent Cas9-based killing experiments.

### In vitro kill-switch evaluation

The kill-switch construct was transformed into BmΔ*vjbR* and recovered on BBA plates supplemented with kanamycin (50 µL/mL) at 37 °C for 3 days. Single colonies were inoculated into 200 µL BBM containing kanamycin (50 µL/mL) in 96-well plates and cultured overnight at 37 °C. Overnight cultures were diluted by transferring 10 µL of culture into 190 µL of fresh BBM containing the antibiotics and grown for an additional 6 h at 30 °C with shaking at 1,000 rpm. Then precultures were diluted to an OD_600_ of 0.1, after which doxycycline was added to a final concentration of 40 ng/mL to induce kill-switch activity. Cultures were incubated for an additional 8 h at 37 °C. Following induction, cultures were diluted to 10^−5^, and 100 µL of each diluted sample was plated on BBA supplemented with kanamycin (50 µL/mL) and doxycycline (40 ng/mL). Plates were incubated at 37 °C for 3 days, and viable cells were quantified by counting colony-forming units (CFUs). Each experiment included three biological replicates and was independently repeated three times.

### In vivo kill-switch evaluation

Mice were intraperitoneally inoculated with 1.07 × 10^9^ colony-forming units (CFU) of the BMΔ*vjbR* harboring either the pBBR1 control plasmid or the pBBR1 plasmid containing a kill-switch construct (n = 2 per group) in a total volume of 100 µL. Mice receiving the BMΔ*vjbR* harboring the empty pBBR1 plasmid served as the vector control, while PBS-injected mice served as negative controls. Three days after inoculation, mice received doxycycline at the indicated dose (0, 20, 25, or 30 mg/kg) by intraperitoneal injection (n=2 per dose).

24 h following doxycycline treatment, mice were euthanized, and livers were collected under aseptic conditions; tissues were weighed and homogenized in 5 mL sterile PBS using the gentleMACs Dissociator (Miltenyi Biotec, Auburn, CA). Tissue homogenates were serially diluted and plated on BBA with kanamycin (50 µg/ml), as appropriate for selection of plasmid-containing bacteria. Plates were incubated at 37°C for 72 h, after which colonies were counted using the Sphere Flash Automatic Colony Counter (Neutec Group Inc., Farmingdale, NY). Bacterial burden was expressed as CFU per gram of tissue. Partial blinding was implemented during bacterial enumeration, whereby samples were identified using animal identification numbers rather than treatment-group designations, and colony counting was performed without knowledge of the corresponding experimental group.

### Ethics approval

Four-to eight-week-old C57BL/6J mice (The Jackson Laboratory, stock no. 000664; Bar Harbor, ME) were housed under specific pathogen-free conditions in a designated Animal Biosafety Level 2 (ABSL-2) animal housing facility at the University of Missouri. Animals were maintained on a 12-h light/dark cycle with ad libitum access to food and water and were provided with appropriate bedding and environmental enrichment, including nesting materials and chewable objects. Sex was not considered an experimental variable in this study, as the primary objective was to evaluate the *in vivo* performance of the engineered kill-switch system rather than sex-dependent differences. All animal procedures were conducted in accordance with the institutional animal care and use guidelines of the University of Missouri and were approved by the University of Missouri Institutional Animal Care and Use Committee (IACUC; protocol no. 44767).

Animals were monitored daily for signs of pain, distress, discomfort, and deterioration in health. Assessments included body weight, body condition, activity level, posture, food and water intake, respiratory status, and general behavior. Indicators of distress included reduced cage exploration, restlessness, reduced mobility, hunched posture, loss of coordination, ruffled fur, reduced grooming, decreased appetite, respiratory distress, and excessive licking, biting, or scratching at injection or procedure sites. Animals did not show any pain and distress and were euthanized at the designated study endpoint, 24 h after doxycycline administration.

Euthanasia was performed by carbon dioxide inhalation, with the flow rate adjusted to displace 30–70% of the chamber volume per minute, followed by cervical dislocation.

### Statistical analyses

Statistical analyses were performed using Microsoft Excel. Data are presented as mean ± standard deviation (SD) unless otherwise indicated. Comparisons between two groups were performed using two-tailed Welch’s t-tests. Each experimental condition included three biological replicates, and all reported *n* values represent biological replicates. A *P* value < 0.05 was considered statistically significant. Statistical significance is denoted as follows: P < 0.05 (*), P < 0.01 (**), P < 0.001 (***), P < 0.0001 (****).

### Novel programs, software, algorithms

Custom computational scripts were developed to perform the model-guided kill-switch design and circuit screening presented in Figure 4. The scripts were used to simulate inducer-dependent Cas9–gRNA complex formation, calculate predicted cell viability, evaluate circuit performance across the design space, and assess the effect of killing sensitivity (K_via_) on circuit behavior. The source code used for these analyses is provided as Supplementary Code 1 accompanying this manuscript.

### Web sites and database references

Candidate gRNAs were evaluated using Benchling (https://benchling.com), which was used to calculate on-target activity and genome-wide off-target scores.

## Data availability Statement

The data supporting the findings of this study are available within the article and its Supporting Information. The source code for all the simulations is available in the Supporting Code.

## Author Contributions

Yeonhwee Jeong: Investigation, Methodology, Validation, Formal analysis, Data curation, Visualization, Writing – original draft, Writing – review & editing.

Hari R. Namboothiri: Investigation, Software, Formal analysis, Visualization, Writing – original draft, Writing – review & editing.

Oluwatola Femi-Olatunji: Investigation, Writing – review & editing.

Paul de Figueiredo: Resources, Funding acquisition, Supervision, Writing – review & editing. Cansu Agca: Supervision, Writing – review & editing.

Chelsea Y. Hu: Conceptualization, Project administration, Funding acquisition, Supervision, Writing – original draft, Writing – review & editing.

## Acknowledgements

This work was supported by the Advanced Research Projects Agency for Health (ARPA-H) under Award Number AY1AX000010. The content is solely the responsibility of the authors and does not necessarily represent the official views of the ARPA-H.

The graphical abstract was partially created using BioRender. Jeong, Y. (2027) https://BioRender.com/2pythn4

We thank Ms. Tracy Mei for technical assistance with plasmid procurement and cloning.

## Supporting Information

Supplementary data is available free of charge at ACS Synthetic Biology online.

## Notes

The authors declare no competing interests.

## Abbreviations

BmΔvjbR: Brucella melitensis ΔvjbR

