## Supplementary Information for "Integrating Genetic Domestication and Model-Guided Kill-Switch Engineering in a Non-Model Bacterial Therapeutic"

**Supplementary Table 1.** Sequences used for construction of the RBS libraries.

| No. | Sequence type | Sequence (5′–3′) |
|---|---|---|
| 1 | ARL library template primer | GAGACTAGTGGGTCTCATACTGAGAAAGA**NNN**GA**NNN**TACAATGAGAGACCTACTAGTA |
| 2 | ARL library reverse primer | CGCTACTAGTAGGTCTCTCA |
| 3 | U3d library template primer | CCGCTTCTAGAGACTAGTGGGTCTCATACTAGAGAA**NNN**GGA**NNN**ATACTAAATGAGAGACCTACTAGTAGCGGCCG |
| 4 | U3d library reverse primer | CGGCCGCTACTAGTAGGTCTCT |

Sequences are shown in the 5′–3′ direction. BsaI recognition sequences are shown in red, and Golden Gate overhangs are shown in blue. Degenerate positions used to generate the RBS libraries are shown in bold. N denotes any nucleotide (A, C, G, or T). All library templates and primers listed in this table were synthesized as single-stranded DNA (ssDNA) oligonucleotides.

**Supplementary Table 2.** Sequences used for construction of the promoter libraries.

| Promoter | Sequence type | Sequence (5′–3′) |
|---|---|---|
| pGroE | Minimal promoter | GCTGTTCGCGCAAAAACGCCAGCCTGAACGGCAATCCTCCTTGACAAAAAAACATTGCGGCTTCTATCTCAAGAACCAGGT |
| | Library template primer | CAAAAACGCCAGCCTGAACGGCAATCCTCCTTG**NNN**AAAAAACATTGCGGCTTCTAT**NNN**AAGAACCAGGTTACTAGAGACCTACTAGTAG |
| | Forward primer | CGCTTCTAGAGACTAGTGGGTCTCAGGAGGCTGTTCGCGCAAAAACGCCAGCCTGAAC |
| | Reverse primer | GCAGCGGCCGCTACTAGTAGGTCTCTAGTAACCTGG |
| pTrcD | Minimal promoter | CAGAAAAAAGATCAAAAAAATTTGACAATTAATCATCCGGCTCGTATAATGTGTGGAATTGT |
| | Library template primer | CTCAGGAGCAGAAAAAAGATCAAAAAAATTTG**NNN**ATTAATCATCCGGCTCGTAT**NNN**GTGTGGAATTGTTACTAGAGACCTAC |
| | Forward primer | CGCTTCTAGAGACTAGTGGGTCTCAGGAGCAGAAAAAAGATC |
| | Reverse primer | CAGCGGCCGCTACTAGTAGGTCTCTAGTAACAATTCC |

Sequences are shown in the 5′–3′ direction. BsaI recognition sequences are shown in red, Golden Gate overhangs are shown in blue, and fragment assembly overlap regions are underlined. Degenerate positions used to generate the promoter libraries are shown in bold. N denotes any nucleotide (A, C, G, or T). Library templates and primers were synthesized as single-stranded DNA (ssDNA) oligonucleotides, while minimal promoter sequences are shown for reference only.

**Supplementary Table 3.** Sequences used for construction of the terminator libraries.

| No. | Sequence type | Sequence (5′–3′) |
|---|---|---|
| 1 | Original T1m | ACTCAAAGCCCGCCGAAAGGCGGGCTTTTCTGTCCTTGAGA |
| 2 | T1m library template primer | GACTAGTGGGTCTCAAGGTACTC**NN**AAGCCCGCCGAAAGGCGGGCTTTT**NN**TGTCCTTG**NNN**GCTTAGAGACCTACTAGTAGCGG |
| 3 | Original T2m | CTAACGCATGAGAAAGCCCCCGGAAGATCACCTTCCGGGGGCTTTTTTATTGCGCCCTTGAGA |
| 4 | T2m library template primer | GACTAGTGGGTCTCAAGGTCTAACGCATGAGA**NN**GCCCCCGGAAGATCACCTTCCGGGGGC**NN**TTTTATTG**NNN**CCTTGAGAGCTTAGAGACCTACTAGTAGCGG |
| 5 | Reverse primer | CCGCTACTAGTAGGTCTC |

Sequences are shown in the 5′–3′ direction. BsaI recognition sites are shown in red, Golden Gate overhangs are shown in blue, and fragment assembly overlap regions are underlined. Degenerate positions used to generate the terminator libraries are shown in bold. N denotes any nucleotide (A, C, G, or T). The same reverse primer was used to construct both the T1m and T2m libraries. Library templates and primers were synthesized as single-stranded DNA (ssDNA) oligonucleotides, while original terminator sequences are shown for reference only.

**Supplementary Table 4.** Candidate gRNA sequences and design scores.

| No. of occurrences in the Bm∆*vjbR* | ON-target score | OFF-target score | gRNA spacer sequence (5′–3′) |
|---|---|---|---|
| 16 | 39 | 99 | GAAAACCGTTTCACACTTTT |
| 12 | 47.9 | 100 | AAAACCGTTTCACACTTTTC |
| 11 | 73.4 | 100 | GGGATCAGAAATCAGTCCAG |
| 7 | 59 | 99 | TCGATCTGATTGAATCAGAT |
| **6** | **66.7** | **100** | **TTTCCAGCAAAAGTGCGAAG** |
| 5 | 55.9 | 100 | AGATCATAGCGCATGCGAGA |

The number of genomic occurrences indicates the number of exact matches of each guide sequence in the Bm∆*vjbR* genome with a compatible NGG PAM site, where N can be any nucleotide. The guide sequence with six genomic occurrences was selected for subsequent experiments and is shown in bold.

**Supplementary Figure 1.** Representative agar plate images from the *in-vitro* kill-switch assay.

| | BmΔ | BmΔ+KS |
|---|---|---|
| (-) Dox | 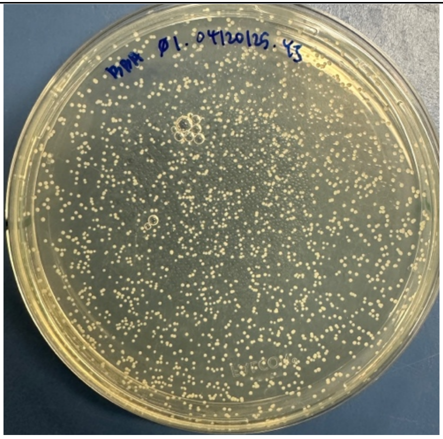 | 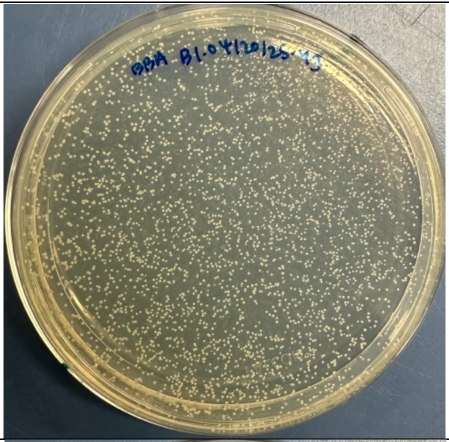 |
| (+) Dox | 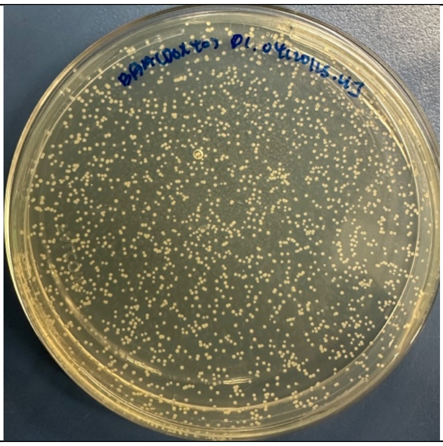 | 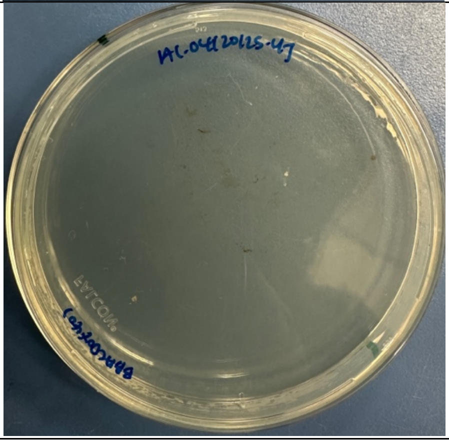 |

**Supplementary Method 1.** Fragment mutagenesis for promoter library construction.

Promoter libraries were generated using a three-primer fragment mutagenesis strategy consisting of a library template primer, a forward primer, and a reverse primer. Each synthetic oligonucleotide was designed to be <120 nt in length. Degenerate nucleotides were introduced at the desired positions within the promoter sequence. Mutations or degenerate positions could be incorporated into the non-annealing regions of any of the three oligonucleotides, provided that sufficient complementary sequence was retained within the annealing regions for primer binding and fragment assembly.

The three oligonucleotides were designed with 18–24 nt overlapping regions to enable reconstruction of the full-length promoter sequence. The resulting assembled promoter fragment was <300 bp in length. PCR amplification was performed using Q5 Hot Start High-Fidelity 2X Master Mix (New England Biolabs, Cat. No. M0494) with an initial denaturation at 98°C for 1 min, followed by 30 cycles of 98°C for 30 s, 57°C for 20 s, and 72°C for 20 s, and a final extension at 72°C for 5 min. Specific oligonucleotide sequences used for the pGroE and pTrcD promoter libraries are provided in Supplementary Table 2.

**Supplementary Method 2.** Dynamic model used for computational screen of kill switch circuit design.

Model Equations

The doxycycline-inducible kill switch circuit was modeled using the following set of ordinary differential equations (ODEs):

$$\frac{dM_{Cas}}{dt} = \beta_{Cas} - (d_m + d_{dil}) \cdot M_{Cas}$$
$$\frac{dP_{cas}}{dt} = k_{tl,Cas} \cdot M_{Cas} - \left(d_{p,Cas} + d_{dil}\right) \cdot P_{Cas} - k_{on} \cdot P_{cas} \cdot G + k_{off} \cdot X$$
$$\frac{dG}{dt} = \beta_G - (d_G + d_{dil}) \cdot G - k_{on} \cdot P_{cas} \cdot G + k_{off} \cdot X$$
$$\frac{dM_{TetR}}{dt} = \beta_{TetR} - (d_m + d_{dil}) \cdot M_{TetR}$$
$$\frac{dP_{TetR}}{dt} = k_{tl,TetR} \cdot M_{TetR} - \left(d_{p,TetR} + d_{dil}\right) \cdot P_{TetR}$$
$$\frac{dX}{dt} = k_{on} \cdot P_{cas} \cdot G - k_{off} \cdot X$$

Detailed descriptions of the species and parameters are included in the tables in this section.

In the screen, $\beta_{Cas}$ and $\beta_G$ are varied between constitutive (strong and weak) and as a $P_{tet}$ promoter. When $\beta_{Cas}$ or $\beta_G$ are modeled as transcription rates under the $P_{tet}$ promoter, their activities are modeled as:

$$\beta = \beta \cdot f_{Ptet}$$

where $\beta$ is either $\beta_{Cas}$ or $\beta_G$, and

$$f_{Ptet} = f_{leak} + \frac{K_{TetR}^n}{K_{TetR}^n + P_{TetR,active}}$$

where $f_{leak}$ is the leaky activity, $K_{TetR}$ is the repression coefficient, $n$ is the Hill coefficient, and $P_{TetR,active}$ represents the active TetR molecules that are not sequestrated by Doxycycline. The concentration of active TetR is given by the equation:

$$P_{TetR,active} = P_{TetR} \cdot \frac{K_{Dox}}{K_{Dox} + Dox}$$

Where $K_{Dox}$ is the repression coefficient and $Dox$ is the concentration of the inducer (Doxycycline).

Model Species

**Supplementary Table 5.** Description of species used in the dynamic model.

| Species | Description |
|---|---|
| $M_{Cas}$ | mRNA of Cas9 |
| $P_{cas}$ | Cas9 protein |
| $G$ | Guide RNA (gRNA) |
| $M_{TetR}$ | mRNA of TetR |
| $P_{TetR}$ | TetR Protein |
| $X$ | Cas9:gRNA complex |

Model Parameter

**Supplementary Table 6.** Description of parameters used in the dynamic model.

| Parameter | Description | Value | Unit |
|---|---|---|---|
| $\beta_{Cas}$ | Maximum transcription rate of Cas9 mRNA | 1 | nM/min |
| $d_m$ | mRNA degradation rate | 1e-1 | 1/min |
| $d_{dil}$ | Dilution due to growth | 6e-3 | 1/min |
| $k_{tl,Cas}$ | Translation rate of Cas9 protein | 1,2 | 1/min |
| $d_{p,Cas}$ | Degradation rate of Cas9 protein | 5e-4 | 1/min |
| $k_{on}$ | Association rate of X | 1e-3 | 1/min.nM |
| $k_{off}$ | Dissociation rate of X | 5e-1 | 1/min |
| $\beta_G$ | Maximum transcription rate of gRNA | 6 | nM/min |
| $d_G$ | gRNA degradation rate | 1e-1 | 1/min |
| $\beta_{TetR}$ | Maximum transcription rate of TetR mRNA | 3 | nM/min |
| $k_{tl,TetR}$ | Translation rate of TetR protein | 1,2 | 1/min |
| $d_{p,TetR}$ | Degradation rate of TetR protein | 5e-4 | 1/min |
| $K_{TetR}$ | Repression coefficient of $P_{tet}$ promoter | 1 | nM |
| $f_{leak}$ | Leaky expression coefficient | 1e-3 | N/A |
| $n$ | Hill coefficient for $P_{tet}$ promoter activity | 1.4 | N/A |
| $K_{Dox}$ | Repression coefficient of active TetR concentration | 2e-3 | ng/mL |

Circuit architecture and transcriptional readthrough model

The circuit architecture comprised three transcriptional units encoding Cas9, gRNA, and TetR. For each unit, both gene order and transcriptional orientation were considered. All permutations of the three components and both possible orientations were generated, while arrangements related by complete reverse complementation were treated as equivalent. This resulted in 24 unique gene-order and orientation arrangements from the ($3! \cdot 2^3$) possible combinations where (3!) represents the possible gene orders, ($2^3$) represents the two transcriptional orientations of each of the three components, and division by two accounts for reverse-complement equivalence. For each arrangement, five Cas9/gRNA promoter configurations and two RBS strengths each for Cas9 and TetR were considered, resulting in ( $24 \times 5 \times 2 \times 2 = 480$ ) unique circuit configurations evaluated in the computational screen.

Transcription from each cassette was first calculated from its assigned promoter activity and maximal transcription-rate coefficient. For cassette $i$, the primary transcription rate was expressed as:

$$\beta_{i,0} = \beta_i \cdot f_i$$

where $\beta_i$ is the maximal transcription-rate and $f_i$ is the corresponding promoter activity. Constitutive promoters were represented by relative activities of 0.1, 1, and 10 for weak, medium, and strong promoters, respectively, whereas $P_{tet}$ activity was determined using the method described in "Model Equations". The relative activities were obtained from the promoter strengths as indicated in the toolbox.

The effect of terminators and cassette arrangement was incorporated through a readthrough model. Each terminator was represented by a readthrough probability, with values of 0.18, 0.02, and 0.01 assigned to weak, medium, and strong terminators, respectively. These values were obtained from the relative terminator strengths as indicated in the toolbox. For a transcript originating from cassette $i$, the transcriptional flux reaching a downstream cassette $j$ was calculated from the product of the readthrough probabilities of all terminators in series as:

$$\beta_{i \to j} = \beta_{i,0} \prod_{k=i}^{j-1} p_k \,,$$

where $p_k$ is the readthrough probability of terminator $k$. When the downstream cassette had the same transcriptional orientation as the originating cassette, the readthrough flux was added to its primary transcription rate. The resulting effective transcription rate for cassette $j$ was therefore:

$$r_j = r_{j,0} + \sum_i r_{i \to j}$$

with the summation including productive same-orientation readthrough contributions. Consequently, the architecture-dependent effect represented in the dynamic model arises primarily from productive transcriptional readthrough between co-oriented cassettes.
